# Gradual sensory maturation promotes abstract representation learning

**DOI:** 10.1101/2025.06.24.661295

**Authors:** Jeonghwan Cheon, Se-Bum Paik

## Abstract

Human infants begin life with limited visual capacities, such as low acuity and poor color sensitivity, due to gradual sensory maturation. In contrast, machine learning models are trained on high-fidelity inputs from the outset, often leading to shortcut learning and overfitting to spurious correlations. Here, we show that early sensory immaturity plays a critical role in shaping bias-resistant, abstract visual representations that conventional models struggle to develop. Using neural network simulations and human psychophysics experiments, we demonstrate that gradual sensory development supports the emergence of robust and generalizable internal representations, reduces reliance on superficial cues, and promotes disentangled representations that enable compositional reconstruction and visual imagination. Comparative analyses of human and model behavior reveal shared patterns of bias resistance and adaptive generalization, including resilience to misleading information. Our findings suggest that gradual sensory maturation is not merely a developmental constraint, but rather a key mechanism that enables abstract representation learning.

**One sentence summary:** Early-stage sensory immaturity guides the development of abstract, bias-resistant representations that enable generalization and compositionality, providing a functional account of gradual sensory maturation.

**Highlights:**

- The essential yet overlooked role of gradual sensory maturation was explored
- Early sensory immaturity promotes abstract representations resistant to shortcut learning
- Emergent representations support compositional reconstruction of novel visual attributes
- Human and model behaviors show similar bias resistance and adaptive generalization

## Introduction

Human perceptual development typically begins with limited sensory capacities, such as low visual acuity and poor chromatic sensitivity, which mature gradually over time ^2–5^. These early limitations have long been regarded as inevitable byproducts of immature sensory and cortical systems, with no specific functional benefit. However, recent evidence suggests that such immature sensory function may play a crucial role in shaping perceptual development ^6^. Studies of late-sighted individuals—patients who gain vision after congenital blindness—provide compelling insights into the role of early sensory experience ^7–10^. Even after surgical restoration, those who begin visual learning with fully mature sensory systems often exhibit persistent deficits in core visual functions, including face recognition ^11,12^, shape processing ^13^, cross-modal integration ^14,15^, and color perception ^16^. Their visual strategies tend to rely on low-level cues, such as local features ^12^ or chromatic contrasts ^16^, rather than high-level, invariant features. These findings suggest that early learning under conditions of sensory immaturity may be crucial for the development of robust visual representations and that its absence can lead to severe cognitive impairments.

Unlike biological systems, machine learning models are typically trained on full-resolution data from the very beginning of the learning process, similar to late-sighted humans. Although these models achieve remarkable performance on standard benchmarks ^17,18^, they remain highly vulnerable to minor perturbations ^19,20^ and shifts in context or environment ^21^.

These failures often stem from shortcut learning ^22^, in which models rely on spurious correlations that are unrelated to the intrinsic structure of the task ^23–26^. Such issues raise concerns about the reliability of machine learning models in real-world applications ^27,28^, where spurious correlations are widespread (Fig. 1) and data distributions often shift unpredictably—conditions humans navigate with far greater resilience.

**Figure 1.**
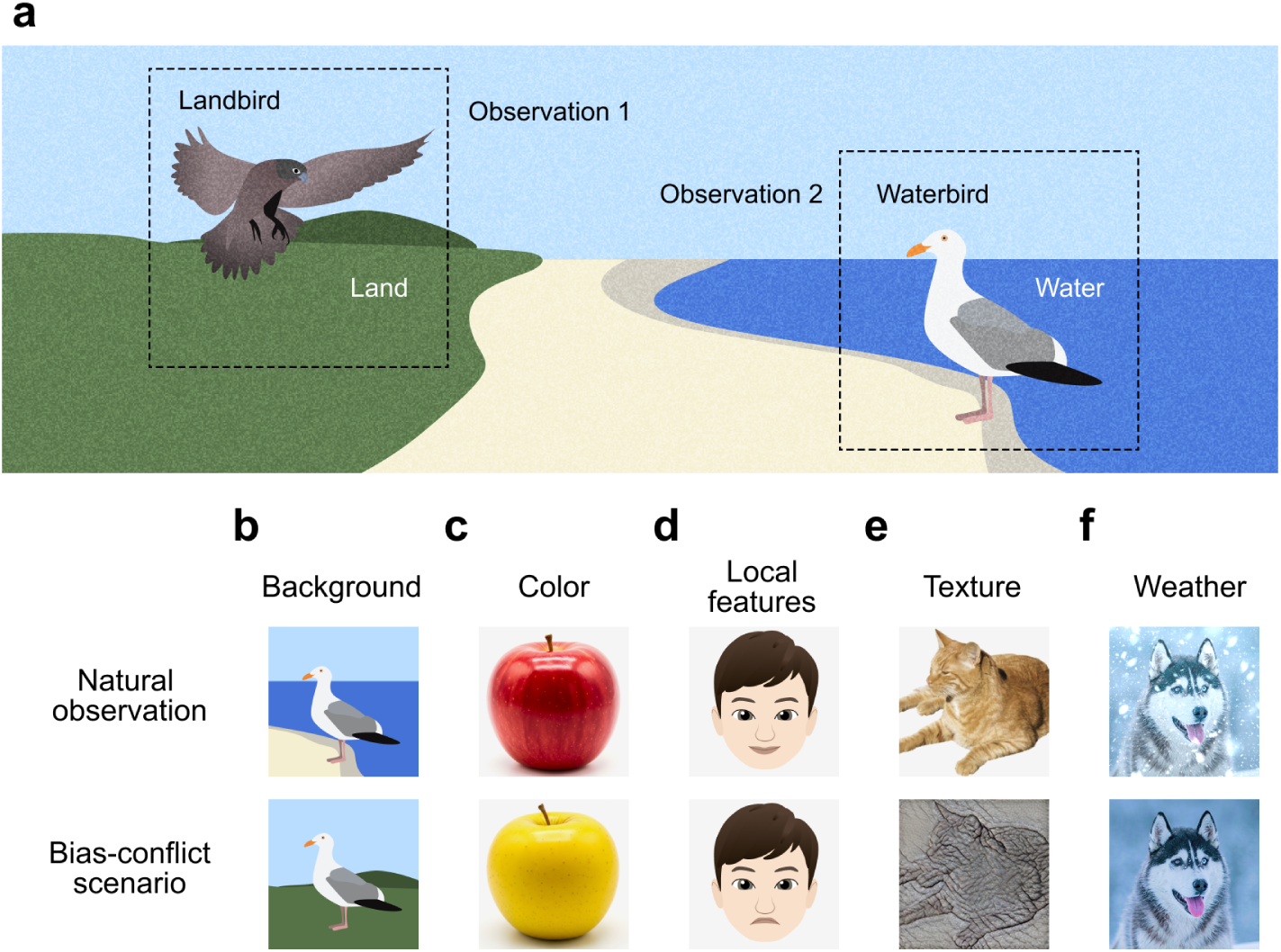
Spurious correlations in natural observations. (a) Illustration of spurious correlations commonly observed in naturalistic data. Landbirds are typically observed against terrestrial backgrounds (Observation 1), whereas waterbirds are more often seen against aquatic backgrounds (Observation 2). (b-d) Examples of different types of spurious correlations and their corresponding bias-conflict, out-of-distribution scenarios, driven by: (b) background, (c) color, (d) local features, (e) texture (image adapted from ^1^), and (f) weather.

Conventional approaches in machine learning often address shortcut learning by adding auxiliary mechanisms, such as extra annotations ^29,30^ or augmentation ^31,32^ to identify and mitigate biases. In contrast, humans develop robust perceptual systems without relying on explicit supervision or auxiliary processing. This contrast raises a fundamental question: Why do artificial neural networks so readily adopt spurious shortcuts while the biological brain acquires resilient perceptual representations? We speculate that the over-reliance on local visual cues observed in late-sighted individuals may mirror the shortcut learning tendencies of machine learning models. In both cases, the absence of early sensory constraints may lead to representational strategies that overfit to superficial features.

Here, we show that gradual sensory maturation can guide perceptual learning and prevent the acquisition of spurious shortcut features, promoting robust and bias-resistant representations. Neural networks initially trained with degraded inputs developed more disentangled and generalizable internal representations that prioritized intrinsic attributes over spurious biases. These models outperformed conventional models on tasks designed to separate intrinsic features from spurious correlations. Their internal representations were more disentangled and robust, supporting both accurate classification and compositional reconstruction of novel stimuli. Comparisons with human behavioral data revealed convergent patterns of bias resistance and adaptive generalization.

Our results suggest that early sensory limitations serve a critical function in shaping robust perceptual representations. Gradual maturation may represent an evolutionarily adapted strategy to avoid shortcut learning, offering a biologically plausible framework for building resilient perceptual systems capable of real-world generalization.

## Results

### Humans rely on intrinsic features, whereas deep neural networks exploit spurious correlations

To investigate how humans and deep neural networks differ in their reliance on intrinsic versus spurious visual features, we designed a perceptual learning paradigm using the Corrupted CIFAR-10 dataset ^31^ (Fig. 2a), in which intrinsic category features (e.g., Airplane, Dog) are systematically paired with bias-inducing corruptions (e.g., Snow, Spatter). In the training set, each category was predominantly associated with a specific corruption type, generating strong spurious correlations (e.g., Snow was consistently applied to Airplane images, and Spatter to Dog images), thereby allowing non-intrinsic, category-independent features to become predictive during training. We refer to these as bias-aligned samples. In contrast, bias-conflict samples — where the category-corruption mapping is broken — serve as critical test cases to assess reliance on intrinsic versus spurious attributes.

**Figure 2.**
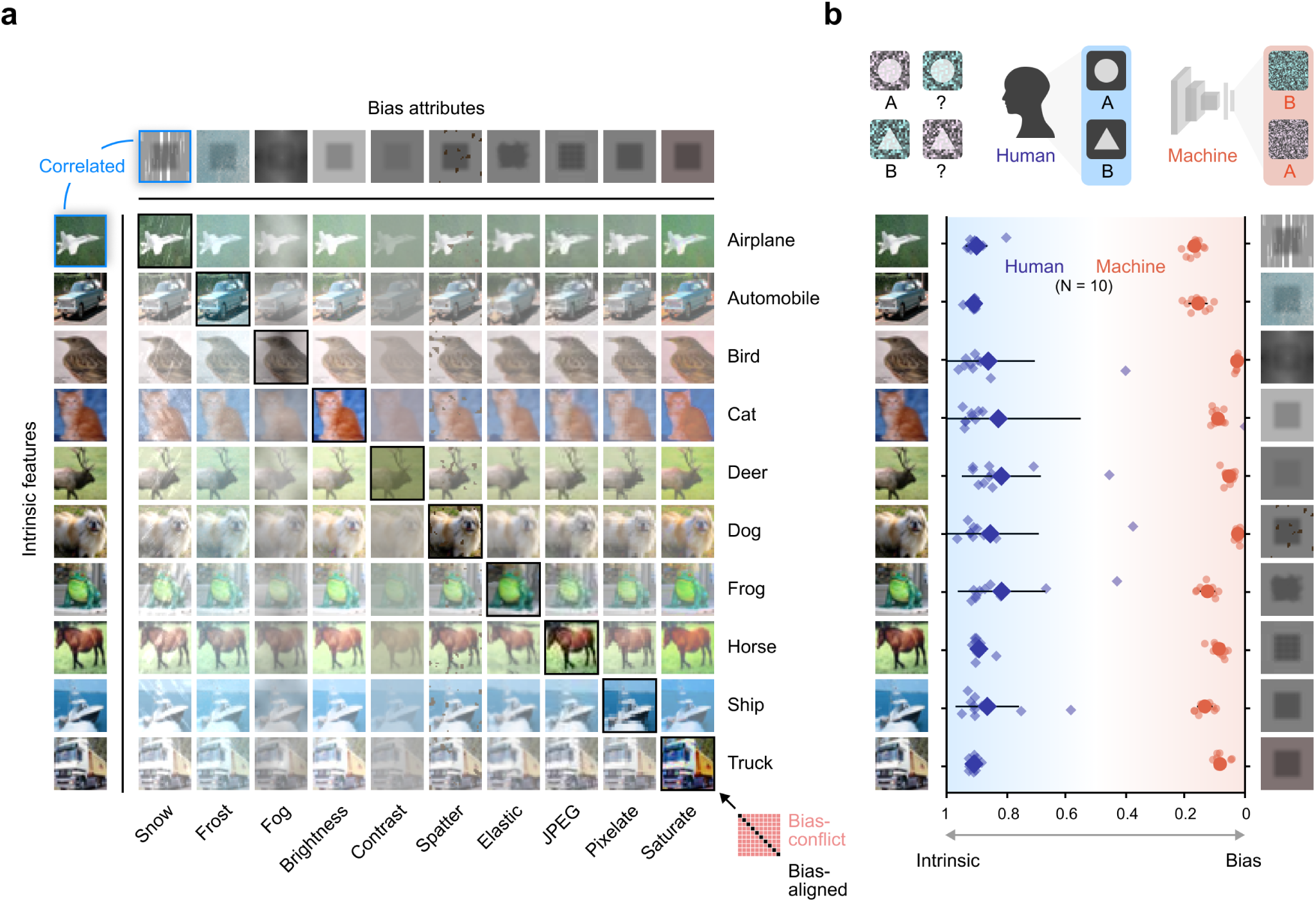
Machine learning models rely on spurious correlations, while humans attend to intrinsic features. (a) Corrupted CIFAR-10 dataset introducing spurious correlations. Each image belongs to one of ten categories (intrinsic features) and is overlaid with one of ten corruption types (bias attributes). In the bias-aligned training set (diagonal), each category consistently co-occurs with a specific corruption (e.g., Airplane – Snow, Automobile – Frost). In the bias-conflict test set, these pairings are disrupted to evaluate generalization. (b) Classification choices on bias-conflict images by machine learning models (orange) and human subjects (purple), both trained only on the biased set (99.5% bias-aligned and 0.5% bias-conflict images). Top: Toy example showing that humans use intrinsic features (e.g., shape), while models rely on spurious ones (e.g., noise color). Bottom: Decision fractions from ten human subjects and models. Values *>* 0.5 reflect reliance on intrinsic features; values *<* 0.5 reflect reliance on bias.

To quantify bias susceptibility, we manipulated the degree of spurious correlation by varying the ratio of bias-aligned to bias-conflict samples. For our main analysis, we used a highly biased training regime containing 99.5% bias-aligned samples and 0.5% bias-conflict samples. Following training, we tested human subjects and neural network models exclusively on bias-conflict samples to determine whether decisions were guided by intrinsic category features or spurious bias cues (Fig. 2b). To quantify this, we introduced a decision fraction metric, defined as the proportion of responses guided by intrinsic attributes relative to those driven by bias attributes. A higher decision fraction indicates greater reliance on intrinsic features, whereas a lower value reflects susceptibility to spurious cues. Human participants (n = 10; 5 male, 5 female; ages 21–26) demonstrated strong reliance on intrinsic features, making perceptual decisions largely independent of the biasing corruptions (Fig. 2b; Supplementary Fig. 1a-d). In contrast, a standard model (ResNet-18^33^) trained on the same biased dataset exhibited near-complete reliance on the spurious bias features, failing to generalize to the bias-conflict samples (Fig. 2b). The difference in decision strategy between humans and the model was statistically significant (Fig. 2b; Human vs. Machine, *n*_subject_ = 10, Wilcoxon rank-sum test, *P <* 10*^−^*^3^; Supplementary Fig. 1e).

To assess the generality of our findings, we replicated the analysis using the Color MNIST dataset (Supplementary Fig. 2a), in which digit identity is spuriously correlated with color in the training set. Both human participants and a machine learning model were trained under these biased conditions (Supplementary Fig. 2b, c). Consistent with the Corrupted CIFAR-10 results, human decisions were reliably guided by the intrinsic feature (digit shape), whereas the machine’s decisions were predominantly driven by the bias attribute (color) (Supplementary Fig. 2d, e; Human vs. Machine, *n*_subject_ = 10, Wilcoxon rank-sum test, *P <* 10*^−^*^3^). These results reinforce the divergence in visual cue utilization between humans and machines: despite achieving similar levels of training accuracy, humans prioritized stable, category-defining features, whereas the machine relied on spurious correlations that hindered generalization beyond the biased training distribution.

### Initial sensory degradation enables debiased learning

Unlike conventional machine learning models that are trained on high-resolution, full-color images from the outset, the biological visual system begins life with limited sensory fidelity (Fig. 3a, left). Human infants initially experience the world with blurry, color-degraded vision, gradually developing acuity and chromatic sensitivity through postnatal sensory and cortical maturation ^3,4^ (Fig. 3a, right). This gradual maturation process may foster robust perceptual learning and mitigate shortcut biases. To test this hypothesis, we simulated the developmental trajectory in artificial neural networks using a staged training paradigm inspired by human vision ^6,12^. Specifically, the “human model” was first trained on blurred, grayscale images, then gradually exposed to high-resolution, full-color inputs (Fig. 3b, Human model). We compared this model to a machine learning model trained on high-quality inputs throughout (Fig. 3b, Machine). Both models were trained on the same strongly biased dataset and achieved similarly low training losses.

**Figure 3.**
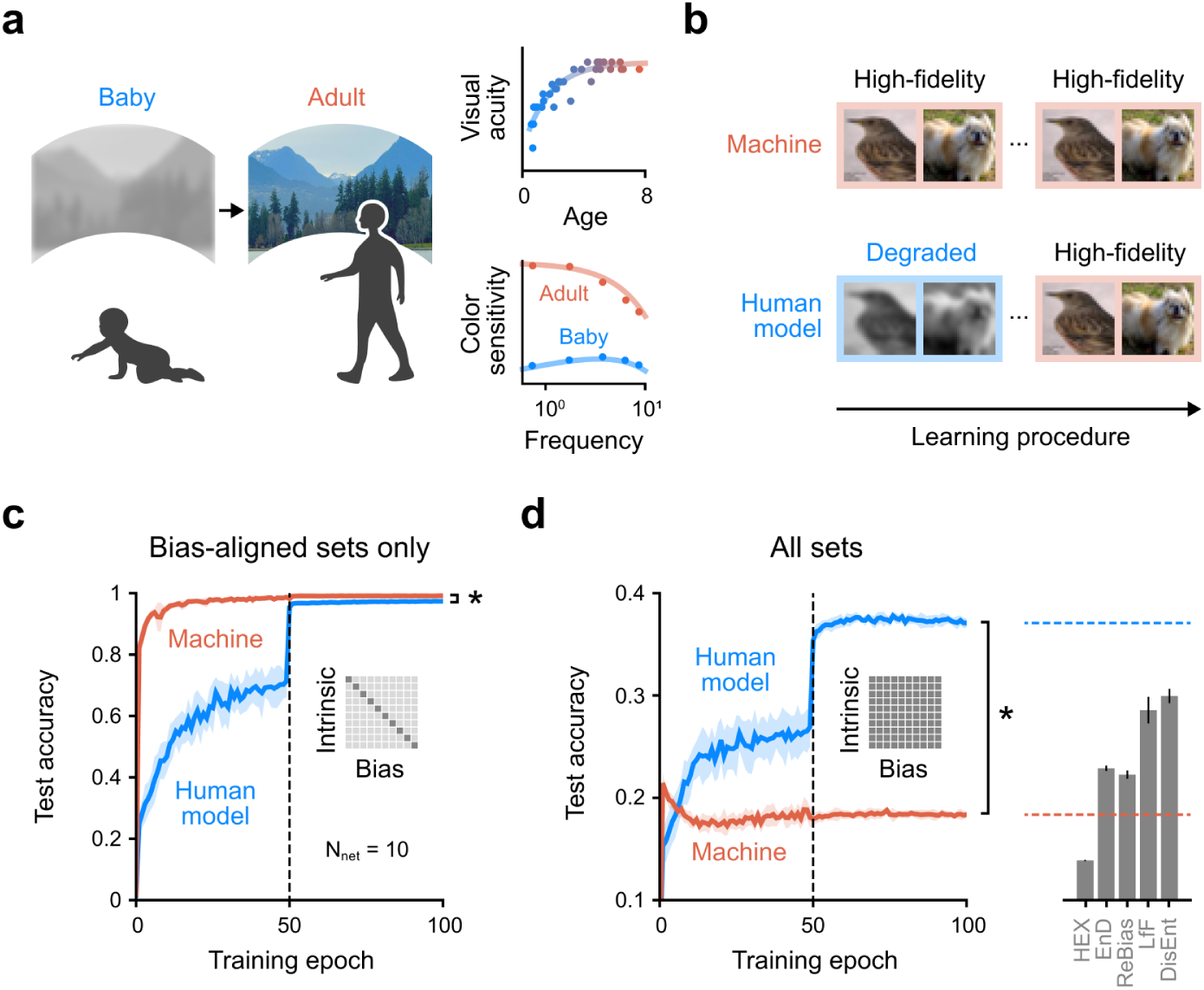
Debiased learning through initial sensory degradation. (a) Learning in biological sensory systems begins with limited capacities that gradually mature. For example, human infants experience blurry and desaturated vision, which refines over time through sensory and cortical development. Right: developmental improvement in visual acuity ^3^ (top) and color sensitivity ^4^ (bottom). (b) Training paradigms for conventional and human-inspired models. The conventional model is trained exclusively on high-quality images (orange). The human-inspired model undergoes gradual sensory maturation — starting with degraded (blurry and grayscale) images, followed by refined ones. Both models are trained on a biased dataset where intrinsic features are spuriously correlated with bias attributes. (c) Test accuracy on a biased test set with spurious correlations preserved. Each data point represents the model’s accuracy on the test set after being trained up to that epoch. The dashed line at epoch 50 denotes the onset of high-fidelity input exposure in the human model’s training. (d) Debiasing performance on an unbiased test set with spurious correlations removed (left) and final benchmark comparisons (right). Gray bars indicate state-of-the-art debiasing methods (HEX ^34^, EnD ^35^, ReBias ^36^, LfF ^31^, DisEnt ^25^). Dashed orange and blue lines mark the final performance of the conventional and human-inspired models, respectively.

To evaluate generalization, we assessed both models on two test sets. On the bias-aligned test set — where spurious correlations between category and bias attributes were preserved — the machine model rapidly achieved high accuracy, whereas the human model improved more gradually (Fig. 3c). By the end of training, both models reached comparable accuracy (Fig. 3c; Human model, 97.33 *±* 0.14%; Machine, 99.13 *±* 0.10%; Human model vs. Machine, *n*_subject_ = 10, Wilcoxon rank-sum test, NS, *P <* 10*^−^*^3^). Notably, on the unbiased test set, the machine model failed to generalize, while the human model improved steadily as it transitioned from degraded to high-quality inputs. By the end of training, the human model significantly outperformed the machine model (Fig. 3d, left; Supplementary Table. 1; Human model, 37.09 *±* 0.44%; Machine, 18.36 *±* 0.32%; Human model vs. Machine, *n*_subject_ = 10, Wilcoxon rank-sum test, *P <* 10*^−^*^3^). The human model also outperformed several state-of-the-art debiasing methods that rely on explicit bias annotations or complex regularization techniques (Fig. 3d, right; HEX ^34^, 13.87 *±* 0.06%; EnD ^35^, 22.89 *±* 0.27%; ReBias ^36^, 22.27 *±* 0.41%; LfF ^31^, 28.57 *±* 1.30%; DisEnt ^25^, 29.95 *±* 0.71%). These findings demonstrate that a simple, biologically inspired training regime can promote robust and bias-resistant representations, outperforming current debiasing algorithms.

One alternative explanation is that the inclusion of degraded inputs alone is sufficient for improving generalization, irrespective of the order in which they are presented. To test this, we evaluated several control models trained with different schedules of sensory degradation (Supplementary Fig. 3). First, models trained exclusively on degraded images exhibited substantially lower performance on both biased and unbiased test sets. Second, in a reverse-curriculum condition, models were initially trained on high-resolution, full-color inputs and then exposed to degraded images. This reverse ordering led to unstable learning dynamics upon the introduction of degraded data, and overall test performance remained poor. These results demonstrate that it is not merely the presence of degraded data, but its early introduction that is critical for robust and debiased learning. Early sensory degradation facilitates resilient internal representations by reducing reliance on shortcut features. To assess the generality of this effect, we tested the same training paradigms on the Color MNIST dataset (Supplementary Fig. 4). Consistent with findings from Corrupted CIFAR-10, both human and machine models achieved high accuracy on the bias-aligned test set. However, on the unbiased test set, the human model again significantly outperformed the machine model, which largely failed to generalize.

We further investigated how different levels of spurious correlation in training data affected model performance. In addition to the primary condition with 99.5% spurious correlation, we evaluated 99%, 98%, and 95% levels (Supplementary Table 1). As expected, unbiased test accuracy increased for both models as the level of spurious correlation decreased. Nevertheless, across all conditions, the human model consistently outperformed the machine model and achieved results comparable with or better than state-of-the-art debiasing methods. These results suggest that early-stage sensory degradation facilitates robust and unbiased learning by limiting shortcut reliance.

### Learning robust representations that capture intrinsic features

Next, we examined the internal representations learned by the human and machine models, focusing on the Color MNIST experiments (Fig. 4; See Supplementary Fig. 5 for results on Corrupted CIFAR-10). We prioritized Color MNIST because color serves as a meaningful and interpretable bias attribute, in contrast to the more arbitrary corruptions used in CIFAR-10. Our aim was to evaluate whether each model’s latent space primarily encoded intrinsic features (digit identity) or spurious biases (digit color). To quantify representational structure, we constructed two latent manifolds for each model by grouping intermediate-layer feature vectors according to either intrinsic or bias labels. At each training epoch, we assessed the linear separability of these manifolds using support vector machine (SVM) classifiers.

**Figure 4.**
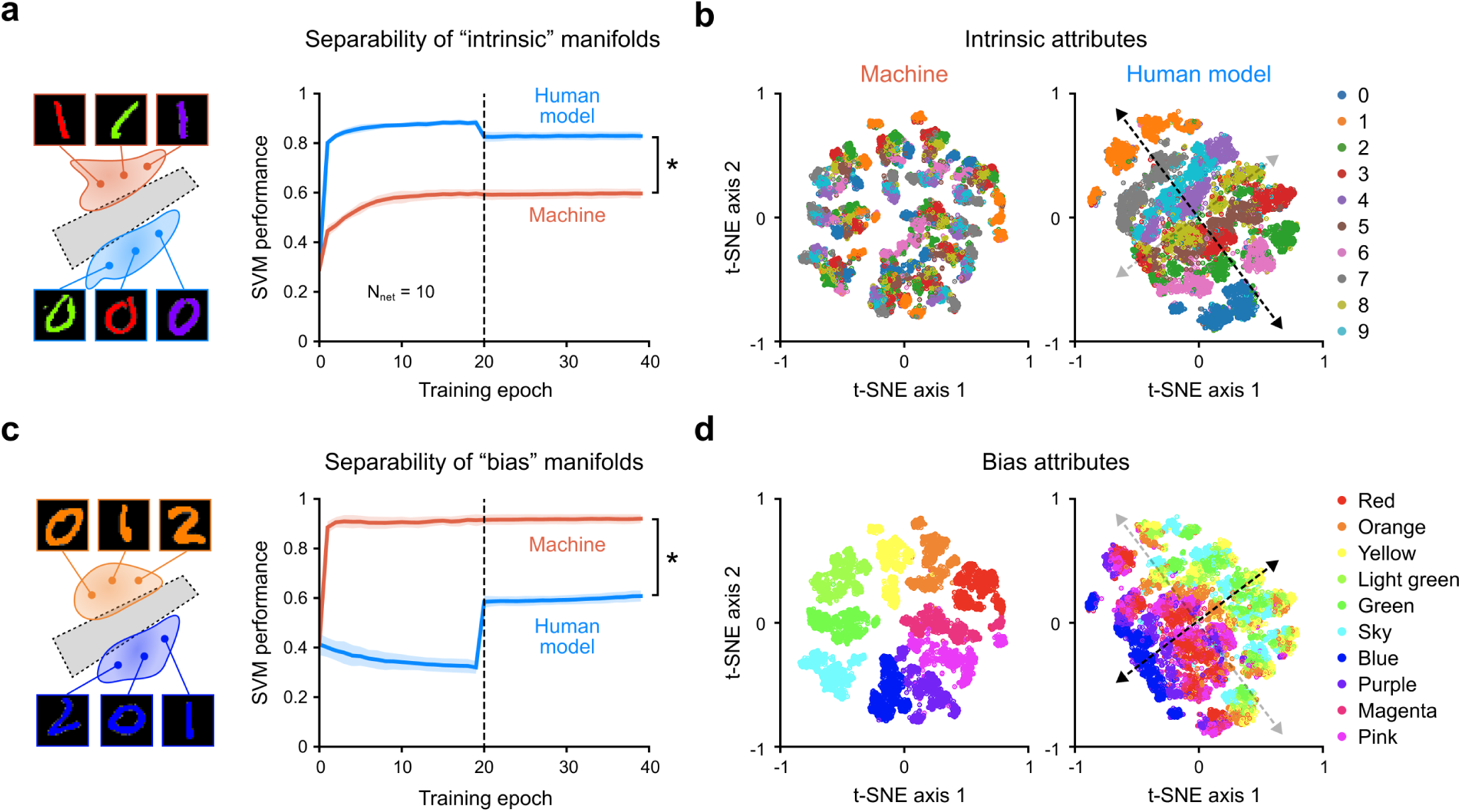
Latent representations in the human-inspired model capture intrinsic attributes over spurious biases. (a) Intrinsic manifold analysis. Latent representations from an intermediate network layer are grouped by intrinsic attributes (e.g., digit identity). Separability is quantified by the average performance of SVM classifiers trained to predict intrinsic categories — higher values indicate stronger intrinsic encoding. Each data point represents the model’s performance after being trained up to that epoch. The dashed line at epoch 20 denotes the onset of high-fidelity input exposure in the human model’s training. (b) Intrinsic structure visualization. Representations from (a) are projected into 2D space using t-distributed stochastic neighbor embedding (t-SNE) and colored by intrinsic attributes. (c) Bias manifold analysis. Same as (a), but representations are grouped by spurious bias attributes (e.g., color), measuring how strongly bias is encoded. (d) Bias structure visualization. Same t-SNE projection as in (b), but colored by bias attributes instead of intrinsic ones.

We first assessed the separability of the *intrinsic* manifold across training (Fig. 4a). The machine model failed to develop representations that separate intrinsic categories (Fig. 4a, Machine), whereas the human model showed a robust increase in linear separability over time (Fig. 4a, Human model vs. Machine, *n*_subject_ = 10, Wilcoxon rank-sum test, *P <* 10*^−^*^3^). This difference was further visualized using t-distributed stochastic neighbor embedding ^37^ (t-SNE), where latent representations from the machine model appeared disorganized with no clear intrinsic clustering (Fig. 4b; Machine), whereas the human model formed distinct clusters aligned with intrinsic labels (Fig. 4b; Human model), largely along a single dominant axis (dashed arrow).

We then evaluated the separability of the *bias* manifold (Fig. 4c). The machine model quickly encoded the bias attribute, achieving high bias separability early in training and maintained this tendency throughout — consistent with its reliance on shortcut cues and high performance on bias-aligned samples. In contrast, the human model suppressed bias encoding, maintaining low bias separability throughout training (Fig. 4c, Human model vs. Machine, *n*_subject_ = 10, Wilcoxon rank-sum test, *P <* 10*^−^*^3^). t-SNE visualizations revealed that the machine model’s latent space clustered strongly by bias features (color) (Fig. 4d; Machine), whereas the human model exhibited a gradual, axis-aligned structure with the bias dimension oriented approximately orthogonal to the intrinsic feature axis (Fig. 4d; Human model, dashed arrow). Thus, the human model forms a disentangled, two-dimensional, map-like latent structure: one axis predominantly encodes intrinsic features, and the other encodes residual bias attributes. Such a structured representation enables the model to capture systematic combinations of relevant and irrelevant features while avoiding entanglement with spurious correlations. We replicated these analyses on Corrupted CIFAR-10 and observed similar patterns (Supplementary Fig. 5). Together, these findings suggest that early-stage sensory degradation promotes the formation of robust and interpretable representations that prioritize intrinsic, task-relevant features while minimizing reliance on spurious, bias-driven signals.

To further test the relationship between internal representations and behavioral outputs, we quantified the representation fraction for each class, measuring the relative contribution of intrinsic features versus bias attributes in the latent space (Supplementary Fig. 6a). Across all classes, the machine model’s representations were dominated by bias-related information, whereas the human model’s representations predominantly reflected intrinsic, category-defining features. We then compared these representational profiles to behavioral decisions by computing the decision fraction — the extent to which perceptual decisions were guided by intrinsic features (Fig. 5). Notably, decision fraction patterns closely mirrored representation fractions: The machine model relied heavily on bias attributes to make decisions (Fig. 5, Machine), while the human model’s decisions were guided by intrinsic features (Fig. 5, Human model). Importantly, these decision patterns closely mirrored those of human subjects (Fig. 5, Human), indicating that models trained with early-stage sensory degradation can effectively reproduce human-like perceptual behavior.

**Figure 5.**
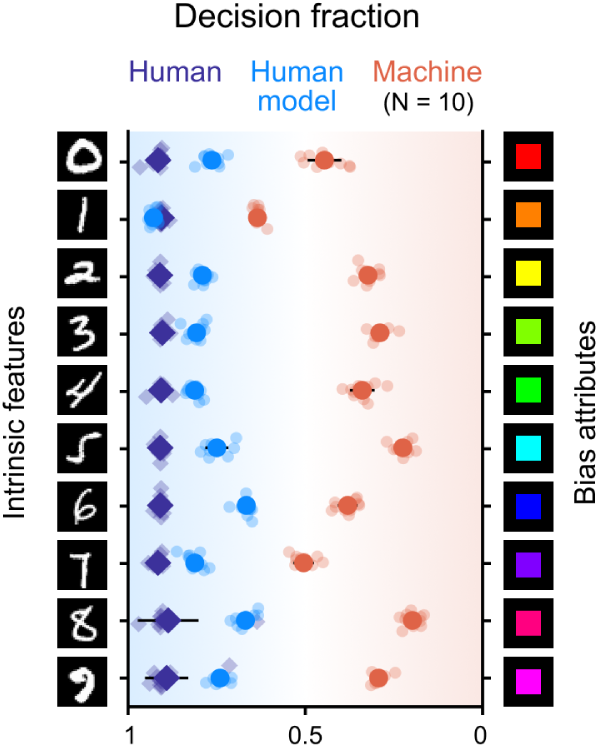
Human-like debiasing in models with early sensory degradation. Decision fraction quantifies the extent to which model decisions rely on intrinsic versus spurious features, ranging from 0 (bias-driven) to 1 (intrinsic-driven). Orange and blue circles represent results from the conventional and human-inspired models, respectively. Purple diamonds indicate human performance from psychophysics experiments.

To directly assess whether the human model’s debiased decisions stemmed from its internal representations, we examined the relationship between representation fraction and decision fraction (Supplementary Fig. 6b). A strong positive correlation was observed between the two (Supplementary Fig. 6b; Regression line, *y* = 1.07*x* + 0.07, *R*^2^ = 0.702; Pearson correlation coefficient, *r* = 0.838, *P <* 10*^−^*^3^), indicating that more intrinsic-feature-dominant representations predict a higher likelihood of intrinsic-feature-based decisions. Notably, the human and machine models formed distinct clusters in this space: the human model exhibited both high intrinsic coding and decision fractions, whereas the machine model consistently relied on spurious features. We observed a similar relationship in the Corrupted CIFAR-10 dataset (Supplementary Fig. 7), reinforcing the conclusion that early sensory degradation shapes internal representations in a way that supports human-like, bias-resistant perceptual decisions.

### Compositional visual reconstruction enabled by abstract, debiased representations

Humans possess a remarkable ability to systematically generalize to novel combinations of visual attributes by leveraging disentangled knowledge, a cognitive capacity known as *systematic compositionality* ^38–41^. For example, someone familiar with “red apple” and “yellow banana” can readily imagine “red banana” or “yellow apple,” even if such combinations are rare or unseen. Prior research has shown that conventional machines struggle with such compositional generalization ^38^. We hypothesized, however, that the abstract representations learned by the human model might enable this form of compositional visual imagination.

To test this, we compared the human and machine models after both were trained on a biased dataset in which specific combinations of intrinsic and bias attributes were overrepresented — for example, “red 0” and “yellow 2.” (Fig. 6a) Their latent representations thus reflected these spurious pairings. We then trained a separate linear visual decoder to reconstruct input stimuli from the latent vectors of each model (Fig. 6a, Decoder). The decoders were evaluated on reconstructing unbiased and compositionally novel combinations — such as “blue 0” or “red 6” — that were rarely seen or absent during training (Fig. 6b). Notably, the decoder trained on the human model’s representation successfully reconstructed novel combinations of features, whereas the decoder trained on the machine model’s representations struggled, particularly under bias-conflict conditions (Fig. 6c, Human model vs. Machine, *n*_subject_ = 10, Wilcoxon rank-sum test, *P <* 10*^−^*^3^). We visualized reconstruction results for all possible combinations of intrinsic and bias attributes (Fig. 6d) and the human model decoder (Fig. 6e). The machine decoder often failed to reconstruct bias-conflict examples, typically preserving the spurious color while misclassifying the shape (Fig. 6f). It frequently reconstructed incorrect digits associated with the spuriously correlated colors, highlighting strong entanglement between intrinsic and bias features. In contrast, the human model decoder consistently reconstructed both shape and color, regardless of whether the combination had been seen during training.

**Figure 6.**
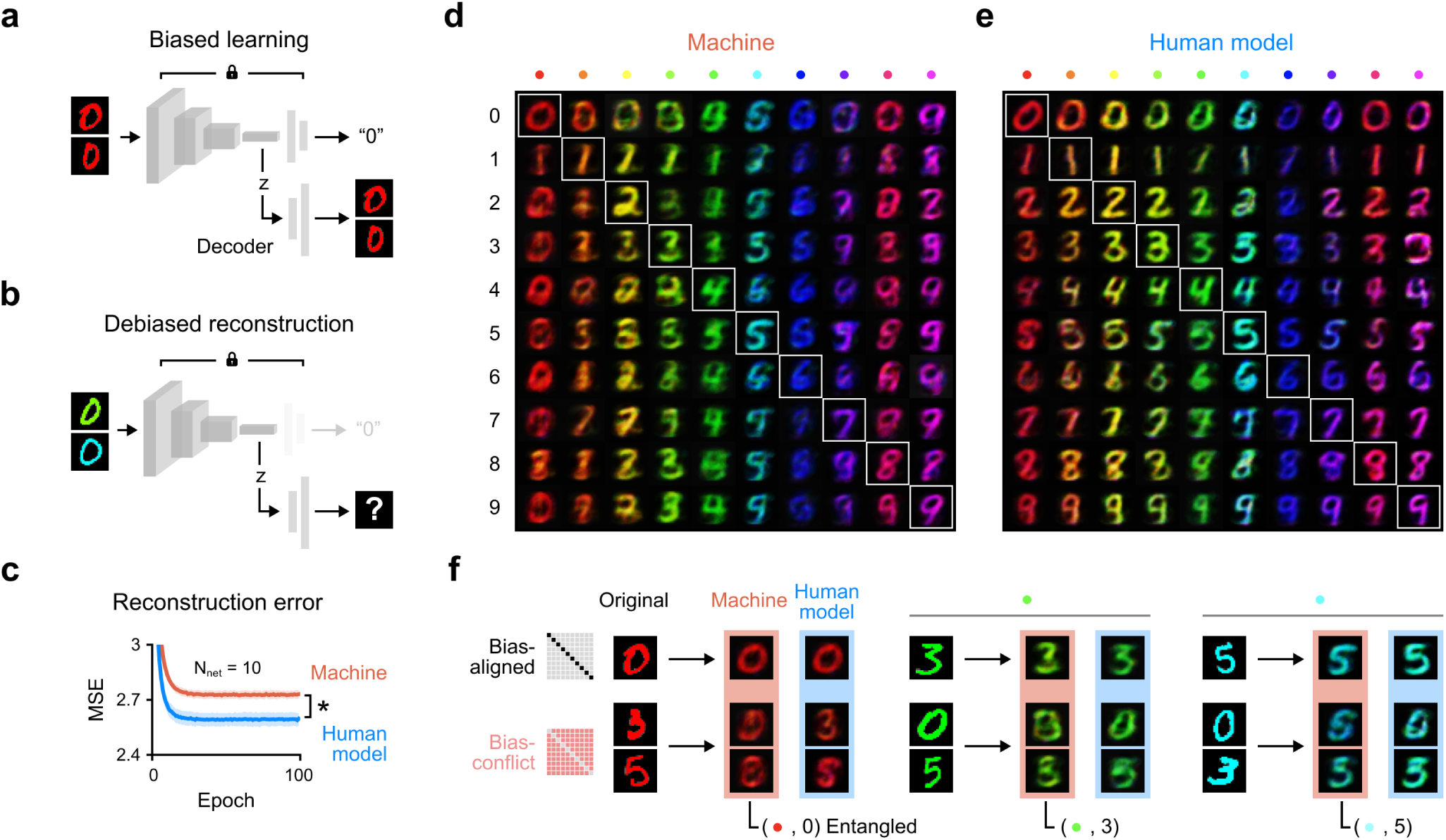
Compositional reconstruction from debiased and disentangled representations. (a) During training, a classifier and a visual decoder are trained on images with strong spurious correlations (e.g., “red 0”, “orange 1”). Latent representations are extracted from an intermediate layer of the classifier, and the decoder learns to reconstruct the original image from these features. (b) The decoder is evaluated on an unbiased dataset containing novel combinations of intrinsic features (digit identity) and bias attributes (color) that were rare or absent during training. (c) Reconstruction accuracy is quantified using mean squared error (MSE) between input images and reconstructions. Orange and blue curves show the performance of the machine and human models, respectively, on the unbiased test set. (d-e) Reconstructed images from the latent representations of the machine model (d) and the human model (e). Rows correspond to intrinsic features (digits) and columns to bias attributes (color). Diagonal entries show reconstructed samples from bias-aligned combinations. (f) Example reconstructions comparing bias-aligned (top row) and bias-conflict (middle and bottom rows) cases. Each original image is shown alongside its reconstructions from both models, illustrating differences in generalization and compositionality.

These results indicate that the human model, shaped by gradual sensory maturation, develops disentangled internal representations that enable compositional visual reconstruction. In contrast to the machine model, the human model flexibly generalizes to novel combinations, suggesting that debiased representations are not only more robust but also inherently more generative. Early-stage sensory limitations, traditionally considered a developmental constraint, may instead serve as a critical inductive bias that supports the emergence of systematic, compositional intelligence. By emulating this biologically grounded developmental process, artificial systems can go beyond data-intensive learning and achieve human-like generalization through principled learning constraints.

## Discussion

Our findings reveal a fundamental principle of perceptual development: early sensory immaturity is not a deficit, but an adaptive mechanism that shapes robust and generalizable visual representations. Gradual sensory maturation plays a critical role in preventing shortcut learning, wherein perceptual decisions rely on spurious, context-specific cues instead of task-relevant, invariant features. This perspective sheds light on why late-sighted individuals often fail to develop invariant object recognition and why conventional machine learning models tend to overfit superficial patterns. By incorporating a model of gradual sensory development, we propose a biologically grounded strategy that mitigates these failures and brings artificial systems into closer alignment with human perceptual behavior.

One particularly striking implication of gradual sensory maturation is its role in enabling compositional visual imagination — the capacity to synthesize novel configurations from learned components. This ability supports core aspects of human intelligence, including generalization to unfamiliar stimuli, flexible categorization, and robust perception under uncertainty. Prior studies have shown that conventional deep learning models often lack systematic compositionality ^38–41^, frequently failing to recombine learned features in ways that generalize beyond the training data. In contrast, we find that models trained with gradually improving sensory input develop internal representations that are not only more robust but also more disentangled, enabling the flexible recombination of learned attributes to reconstruct novel inputs. These results suggest that early sensory limitations may scaffold the development of high-level, generative perceptual capacities. Notably, disentangled and abstract representations — supporting adaptive generalization and imagination — are observed in higher-order regions of the human brain ^42–46^. Our research provides a developmental account for the emergence of such representations grounded in an initial sensory bottleneck.

Recent research has revisited early perceptual development in children to explore how biologically inspired processes can support robust representation learning in artificial systems ^6,12–16,47^. Computational studies, in particular, have shown that early exposure to degraded visual input can enhance generalization and robustness across various domains. For example, initial visual degradation has been found to promote broader receptive fields, improving face recognition performance without relying on local features ^12^. Training with blurry images increases the robustness of the model and shifts the biases of the network from texture to shape, in closer alignment with human perception ^47^. Likewise, early color degradation can lead to more stable and resilient chromatic representations ^16^.

Although these findings emphasize the benefits of developmental constraints, they have often been treated as disconnected effects. Here, we offer a unifying framework by interpreting them as instances of shortcut learning, where both artificial networks and late-sighted individuals exploit spurious, low-level cues such as local facial details, textures, or color signals. From this perspective, early sensory limitations help suppress shortcut biases and promote representations based on intrinsic, task-relevant features. This interpretation connects the well-documented shortcut-learning problem in machine learning ^22^ with the cognitive impairments observed in late-sighted individuals ^10,12,16^. By reframing these developmental effects in terms of shortcut avoidance, our framework offers a theoretical basis for designing more robust and human-aligned machine learning systems, as well as new insights into potential strategies for remediating perceptual deficits in atypical developmental trajectories.

This process of perceptual development, grounded in early sensory limitations, follows a trajectory in which both the quantity and complexity of sensory input gradually increase over time. In the earliest stage, the biological brain processes brief and simplified input, gradually building toward more detailed and structured representations. A comparable strategy in machine learning is curriculum learning, where tasks are presented in order of increasing difficulty to promote generalization more effectively than näıve training ^48^. From this perspective, gradual sensory maturation can be viewed as a biologically embedded and evolutionarily adapted form of curriculum learning. Unlike conventional curriculum learning in artificial systems, which requires carefully engineered task sequences or data schedules, developmental curriculum learning emerges naturally from early sensory constraints, without the need for externally designed instruction.

Our recent studies draw direct inspiration from key developmental milestones in the biological brain ^49–51^. During early prenatal stages, spontaneous neural activity in the sensory cortex resembles random noise ^52,53^. Using this phase, we introduced a biologically plausible noise pretraining regime and showed that it improves learning speed, accuracy, and robustness — even under local learning rules ^50,54^. We further demonstrated that such pretraining improves uncertainty calibration, a computational analog of “metacognition” — the ability to evaluate the confidence of one’s own decisions ^51^. Extending this line of work, we modeled the postnatal period, during which infants experience degraded sensory inputs that gradually refine over time. We found that training with initially degraded inputs not only prevents shortcut learning on spurious correlations but also supports more human-aligned generalization. Together, these approaches inspired by developmental neuroscience address critical challenges in machine learning. They illustrate the broader promise of NeuroAI ^55–60^, demonstrating how biologically grounded strategies can help align artificial learning systems more closely with the structure and dynamics of human cognition.

In conclusion, our findings offer a biologically motivated and computationally effective solution to a long-standing limitation of machine learning models. More broadly, these results highlight the functional role of early sensory constraints in shaping robust perceptual development and suggest that principles of neurodevelopment can inform the design of more resilient and human-aligned machine learning systems.

## Methods

### Construction of image datasets with spurious correlations

We used benchmark datasets with spuriously correlated biases to investigate the learning behavior and perceptual decisions of both machine learning models and human subjects. Each dataset consisted of intrinsic features and bias attributes that were artificially correlated in the training data. To evaluate generalization performance, we employed two types of test sets. The bias-aligned test set included only samples that preserved the spurious correlations seen during training. In contrast, the unbiased test set contained images representing all possible combinations of intrinsic and bias attributes with equal sampling, resulting in a large proportion of bias-conflict samples.

For object recognition, we used the Corrupted CIFAR-10 dataset—a modified version of CIFAR-10 in which spurious correlations are introduced via systematic visual corruptions ^31^. The original CIFAR-10 consists of ten object and animal categories: Airplane, Automobile, Bird, Cat, Deer, Dog, Frog, Horse, Ship, and Truck, which serve as the intrinsic attributes ^61^. We applied ten corruption types: Snow, Frost, Fog, Brightness, Contrast, Spatter, Elastic, JPEG compression, Pixelate, and Saturate ^62^ as bias attributes. Each intrinsic attribute is spuriously correlated with one bias attribute; for example, the Snow corruption is applied exclusively to Airplane images. Samples exhibiting this spurious correlation are defined as bias-aligned samples, while all other combinations are considered bias-conflict samples. For the training dataset, we created four variants with different degrees of spurious correlation: 99.5%, 99%, 98%, and 95%. For instance, a 99.5% correlation ratio means that 99.5% of the samples exhibit the spurious correlation, while only 0.5% are bias-conflict examples. For our main results, we used the dataset with a 99.5% spurious correlation.

To assess digit classification, we used the Color MNIST dataset, a modified version of MNIST in which digit identities (intrinsic features) are spuriously correlated with color (bias attributes) ^31^. The original MNIST dataset contains handwritten digits from 0 to 9^63^. We applied ten distinct colors — Red, Orange, Yellow, Light Green, Green, Sky, Blue, Purple, Magenta, and Pink — such that each digit was predominantly associated with a specific color (e.g., most images of the digit “0” were colored red). As with Corrupted CIFAR-10, we constructed four versions of this dataset (99.5%, 99%, 98%, and 95% spurious correlation) and employed the same training and testing protocols. All primary results were based on the 99.5% version.

### Neural network model architectures

We employed convolutional neural networks and multilayer feedforward networks as representative conventional machine learning models to train on spuriously correlated image datasets. For the Corrupted CIFAR-10 datasets, we used ResNet-18^33^, following prior studies that utilized the same benchmark ^25,26,31^. ResNet-18 consists of a feature extraction network and a classification network. The feature extraction network begins with a convolutional layer, followed by batch normalization, a rectified linear unit (ReLU) activation, and a max pooling layer. This is followed by four groups of residual blocks, each containing two residual blocks with convolutional layers and identity skip connections to facilitate effective gradient flow. The classification network includes a global average pooling layer that aggregates spatial features into a compact vector and a fully connected layer that produces the final class predictions. For the Color MNIST experiment, we used a four-layer feedforward network with a hidden layer size of 100-100-32. This network consists of fully connected layers, followed by ReLU activations, which ultimately output the final class predictions.

### Training Procedures: Conventional vs. Gradual sensory maturation

To compare conventional machine learning models with human-inspired models that incorporate gradual sensory maturation, we examined two distinct training procedures. In the standard approach, models were trained on high-resolution, full-color images throughout the entire training process. In contrast, the human-inspired approach began with an initial phase of degraded inputs, followed by training on high-quality images. To simplify the process, networks were trained on degraded data for the first half of the epochs. During this degradation phase, input images were transformed using Gaussian blurring and converted to grayscale. For the Corrupted CIFAR-10 experiments, we applied a Gaussian filter with a kernel size of 7 and a standard deviation ranging from 0.5 to 2. For the Color MNIST experiment, we used a kernel size of 3 with the same range of standard deviations. All image transformations were implemented using PyTorch transformation functions. The second half of the training was conducted using the original, unaltered images.

We employed standard neural network training procedures using the Adam optimizer with *β*_1_ = 0.9 and *β*_2_ = 0.999. For both the Corrupted CIFAR-10 and Color MNIST experiments, the initial learning rate was set to 0.0001, and a batch size of 128 was used. Networks were trained for 100 epochs in the Corrupted CIFAR-10 experiment and for 40 epochs in the Color MNIST experiment. To facilitate convergence, a learning-rate decay (*γ* = 0.1) was applied at the midpoint of training. All input images were normalized using the mean and standard deviation calculated from the training set. For the Corrupted CIFAR-10, random horizontal flipping and random cropping were applied during training as data augmentation techniques to mitigate overfitting. No data augmentation was used during testing. Training hyperparameters were carefully tuned to ensure improved performance and stable learning while maintaining fair comparisons between the conventional and human-inspired models. Notably, the results remained consistent and robust across variations in these hyperparameters.

### Human psychophysics experiments on bias sensitivity

To evaluate human perceptual decisions on spuriously correlated images, we conducted psychophysical experiments with ten human subjects (5 males, 5 females; ages 21–26) with normal or corrected-to-normal vision. Written informed consent was obtained in accordance with protocols approved by the Institutional Review Board (IRB) of KAIST (KH2021-013). All procedures adhered to the approved ethical guidelines.

The experiment consisted of two sessions corresponding to the Corrupted CIFAR-10 and Color MNIST datasets, each comprising a learning phase followed by a test phase. In the learning phase, participants completed 40 trials. Each trial began with a 0.5-second fixation cross, after which ten example stimuli from different object categories were presented in random order along the top of the screen. Below, ten response boxes were displayed. Participants were asked to drag and drop each image into the box they believed matched its category. Participants have a limited time to complete their responses (20 seconds for the Corrupted CIFAR-10; 15 seconds for Color MNIST). Immediate feedback was provided (“O” for correct, “X” for incorrect), allowing participants to learn the category-label associations. Stimuli during this phase were drawn from a biased dataset with 99.5% bias-aligned examples, replicating the conditions used in neural network training. The test phase also comprised 40 trials using the same presentation and response format but without feedback. Test stimuli were drawn from an unbiased dataset in which intrinsic and bias attributes were independent. Participants’ responses in this phase were used to assess whether categorization relied on intrinsic features rather than spurious cues. Practice sessions were conducted before the main experiment to familiarize participants with the task. Additional controls—including randomized trial order and standardized viewing conditions—were implemented to ensure experimental rigor and minimize confounds commonly encountered in psychophysical testing.

### Quantifying bias in perceptual decision-making

To evaluate perceptual decision-making on images containing spuriously correlated features, we measured responses on a bias-conflict test set. In these images, intrinsic attributes (e.g., object or digit identity) and bias attributes (e.g., corruption type or color) were deliberately mismatched. This design enabled us to quantify the extent to which decisions were driven by intrinsic features versus bias attributes. For each conflict image, the decision was recorded as being consistent with either the intrinsic attribute or the bias attribute. Following a procedure similar to the shape–texture bias paradigm ^1^, we defined the decision fraction (DF) as

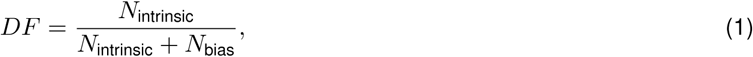

where *N*_intrinsic_ is the number of trials in which the decision matched the intrinsic features, and *N*_bias_ is the number of trials in which the decision aligned with the bias attribute. A DF value greater than 0.5 indicates a preference for intrinsic features, whereas a value below 0.5 suggests stronger reliance on spurious biases. The metric was applied consistently across both machine learning models and human subjects to assess their susceptibility to bias in decision-making.

### Manifold analysis of latent representations

To characterize the latent representations, we conducted manifold analysis on feature vectors extracted from the penultimate layer of the network after applying the activation function. Two types of manifolds were defined using the same hidden representations: intrinsic manifolds, in which vectors were grouped according to intrinsic labels (e.g., digit or object class), and bias manifolds, grouped by spurious attributes (e.g., corruption type or color). We first applied principal component analysis (PCA) to reduce dimensionality, projecting each manifold onto its ten leading principal components. To quantify separability, we used support vector machines (SVMs). Specifically, since there were ten categories, we trained ten one-versus-rest SVM classifiers, each distinguishing one manifold from all others. The separability score was defined as the mean classification accuracy across the ten SVMs. This approach allowed for a direct, quantitative comparison of how well intrinsic versus bias features were represented and separated in the model’s latent space.

### Visualization of latent representation

To visualize the latent representations of our neural network models, we extracted the hidden activations from the penultimate layer after applying the activation function. To capture the low-dimensional structure of these representations, we applied t-distributed stochastic neighbor embedding (t-SNE) analysis ^37^. t-SNE projects the high-dimensional latent space onto a two-dimensional plane while preserving local relationships between samples. We visualized the resulting embeddings in two ways: (1) by coloring each point according to its intrinsic label (e.g., digit or object class) and (2) by coloring each point according to its bias attribute (e.g., corruption type or color). These visualizations allow us to assess how well the latent space distinguishes between intrinsic and bias features, offering a comprehensive view of the structure and separability of the internal representations.

### Quantifying latent representations

To quantify the extent to which latent representations encode intrinsic attributes versus bias attributes, we introduced the representation fraction (RF), a metric analogous to the decision fraction but applied to the latent representations from the network’s penultimate layer. To assess the quality of clustering, we used the silhouette coefficient, defined for each sample *i* as

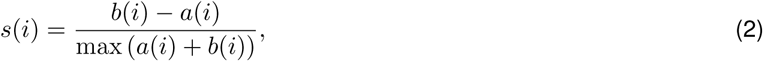

where *a*(*i*) is the mean intra-cluster distance for point *i*, and *b*(*i*) is the mean distance to the nearest cluster. The silhouette coefficient ranges from -1 to 1: Positive values indicate that the sample is well matched to its own cluster and poorly matched to others, while negative values indicate the opposite.

We computed two average silhouette coefficients: *A*_intrinsic_, the average silhouette coefficient when samples are clustered by intrinsic labels, and *A*_bias_, the average when clustered by bias attributes. The representation fraction (RF) was then calculated as

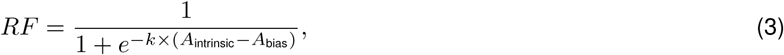

where *k* is a scaling parameter controlling the slope of the sigmoid function. We used *k* = 5 to match the scale of the RF to that of the decision fraction. An RF greater than 0.5 indicates that intrinsic attributes are more distinctly encoded (i.e., more separable) than bias attributes in the latent space, whereas a value below 0.5 suggests the dominance of bias-driven encoding. This metric provides a quantitative measure of representational bias in the internal feature space of the model.

### Visual reconstruction to probe debiased and disentangled representations

To evaluate whether the latent representations are both debiased and disentangled, we designed a visual reconstruction task based on features extracted from the latent representations. A linear visual decoder was trained to reconstruct the input image from the latent representation:

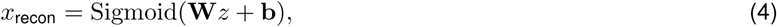

where *z* is the latent feature vector, **W** and **b** denote the decoder’s weights and bias, respectively. The decoder consists of a single linear layer followed by a Sigmoid activation and outputs an image of size 28 *×* 28 *×* 3. Latent features were obtained from a previously trained classifier, and the classifier’s weights were kept fixed during the reconstruction phase. Both the classifier and decoder were trained on biased datasets in which intrinsic and bias features were strongly correlated.

During evaluation, the decoder was tested on an unbiased dataset, composed largely of bias-conflict samples — stimuli featuring combinations of intrinsic and bias attributes that were rare or absent during training. For example, while the model may have been trained primarily on “red 0” digits, the test set included “0” digits in orange or green. This setup allowed us to assess the generalization capacity of the model’s latent representations: whether they encode intrinsic characteristics independently of spurious bias attributes. Successful reconstruction of such novel combinations would indicate that the model has learned abstract, compositional representations that are not tied to spurious correlations.

### Statistical analysis

All statistical details, including sample sizes, exact P values, and the specific statistical methods used, are reported in the main text or the corresponding figure legends.

## Data availability

The datasets used in this study are publicly available. Color MNIST and Corrupted CIFAR-10 are available in https://github.com/kakaoenterprise/Learning-Debiased-Disentangled. These spurious correlated image sets were generated by synthesizing bias attributes from the original MNIST (http://yann.lecun.com/exdb/mnist/) and CIFAR-10 (https://www.cs.toronto.edu/~kriz/cifar.html) datasets. The codes for generating Color MNIST and Corrupted CIFAR-10 are available at https://github.com/alinlab/LfF, a repository from previous research introducing these datasets.

## Acknowledgements

This work was supported by the National Research Foundation of Korea (NRF) grants (NRF2022R1A2C3008991 to S.P.) and by the Singularity Professor Research Project of KAIST (to S.P.).

## Author contributions

Conception and design: J.C. Development of the methodology: J.C. and S.P. Funding acquisition: S.P. Collection of data: J.C. Formal analyses: J.C. and S.P. Writing the original draft: J.C. and S.P. Writing, reviewing and/or revising the manuscript: J.C. and S.P. Study supervision: S.P.

## Supplementary Materials

**Supplementary Fig. 1.**
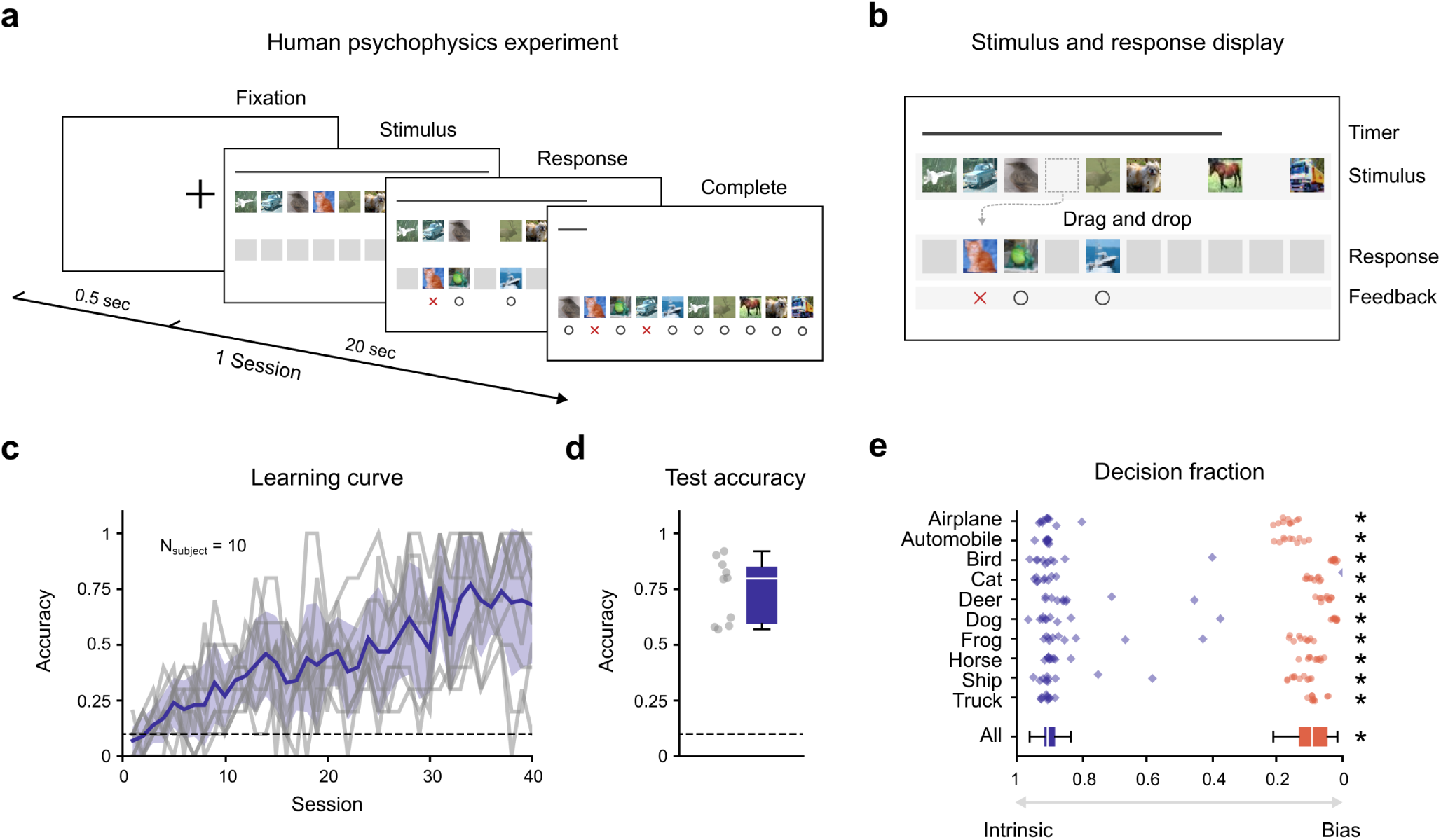
Psychophysics experiment evaluating human learning under spurious correlations. (a) Experimental design. Each session begins with a 0.5-second fixation period, followed by the stimulus display. Participants have a limited time to complete their responses (20 seconds for the Corrupted CIFAR-10; 15 seconds for Color MNIST) by dragging all ten stimuli into their corresponding response boxes. (b) Stimulus and response interface. Ten stimuli are shown in randomized positions, with a time bar displayed at the top of the screen. During training, participants learn the correct category-box mapping through trial and error with immediate feedback (“O” for correct, “X” for incorrect). No feedback is provided during the test phase. (c) Learning curves for ten human subjects (gray lines) and the group average (purple line). (d) Test performance across 40 trials per participant (400 responses total). (e) Decision fractions for ten human subjects (purple) and the machine model (orange) after training on the ten Corrupted CIFAR-10 categories. The decision fraction reflects reliance on intrinsic versus bias features: values *<* 0.5 indicate bias-driven decisions, and values *>* 0.5 indicate intrinsic-driven decisions. Human participants relied significantly more on intrinsic features than the machine model (Human vs. Machine, *n*_subject_ = 10, Wilcoxon rank-sum test, *P <* 10*^−^*^3^). This tendency was consistent across all intrinsic–bias pairs (Human vs. Machine, *n*_subject_ = 10, Wilcoxon rank-sum test; Airplane - Snow, Automobile - Frost, Bird - Fog, Cat - Brightness, Deer - Contrast, Dog - Spatter, Frog - Elastic, Horse - JPEG, Ship - Pixelate, Truck - Saturate, *P <* 10*^−^*^3^).

**Supplementary Fig. 2.**
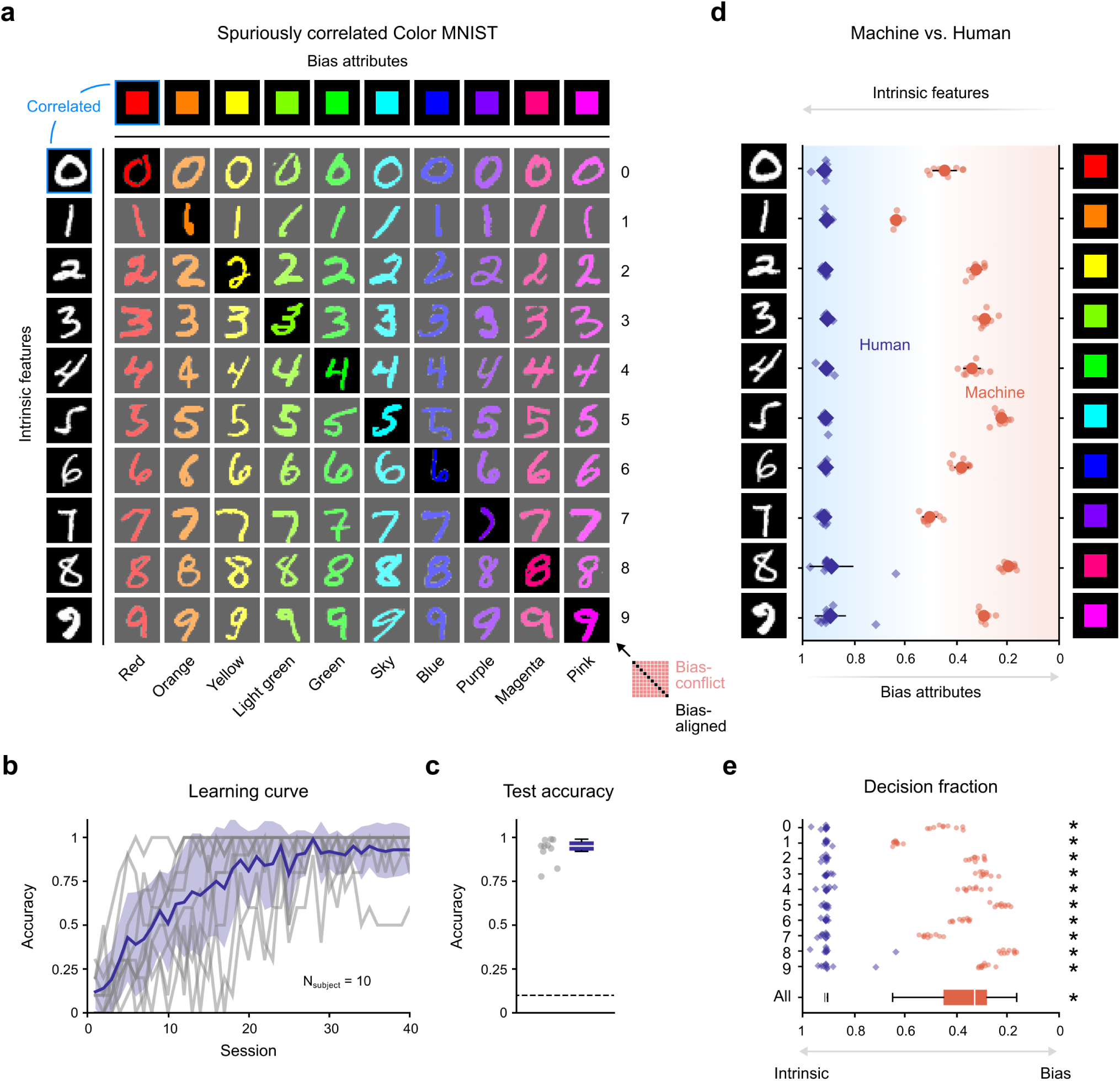
Color MNIST dataset and human learning under spurious correlations. (a) The Color MNIST dataset was used to introduce spurious correlations between digit identity (intrinsic feature) and color (bias attribute). In the bias-aligned training set, each digit is consistently paired with a specific color (e.g., “0” with Red, “1” with Orange), forming strong but artificial correlations (shown on the diagonal). The bias-conflict test set includes mismatched combinations to evaluate generalization beyond the learned bias. (b) Learning curves for ten human subjects (gray lines) and their average learning curve (purple line). (c) Test performance across 40 test sessions per participant (400 total responses). (d) Decision fractions on bias-conflict test images for machine learning models (orange) and human subjects (purple), both trained only on the bias-aligned dataset. (e) Decision fractions by digit–color pair. Human participants show significantly greater reliance on intrinsic features compared to machine models (Human vs. Machine, *n*_subject_ = 10, Wilcoxon rank-sum test, *P <* 10*^−^*^3^). This tendency held consistently across all digit-color pairs (Human vs. Machine, *n*_subject_ = 10, Wilcoxon rank-sum test; “0” - Red, “1” - Orange, “2” - Yellow, “3” - Light green, “4” - Green, “5” - Sky, “6” - Blue, “7” - Purple, “8” - Magenta, “9” - Pink, *P <* 10*^−^*^3^).

**Supplementary Fig. 3.**
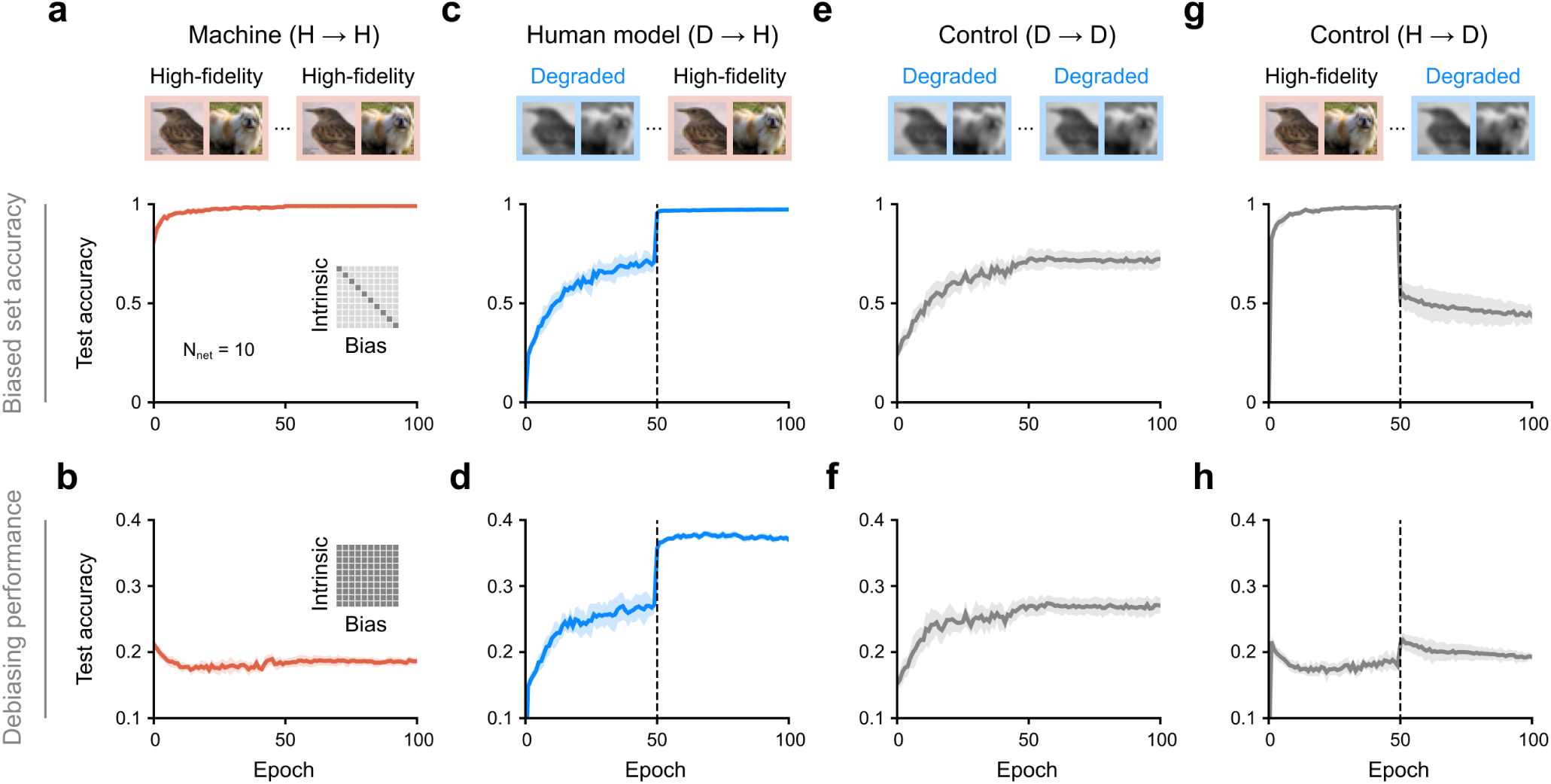
Importance of training order for degraded versus high-fidelity inputs. (a-b) Conventional models trained only on high-fidelity (refined) images throughout training (High-fidelity to High-fidelity). Test accuracy is plotted over training epochs on the biased test set (a) and unbiased test set (b). (c-d) Human-model training procedure (Degraded to High-fidelity): Training begins with degraded images (blurred and grayscale) for the first half and switches to refined images at epoch 50 (dashed line). (e-f) Control models trained exclusively on degraded images for the entire training period (Degraded to Degraded). (g-h) Reversed curriculum control (High-fidelity to Degraded): Training starts with refined images and switches to degraded inputs at epoch 50 (dashed line).

**Supplementary Fig. 4.**
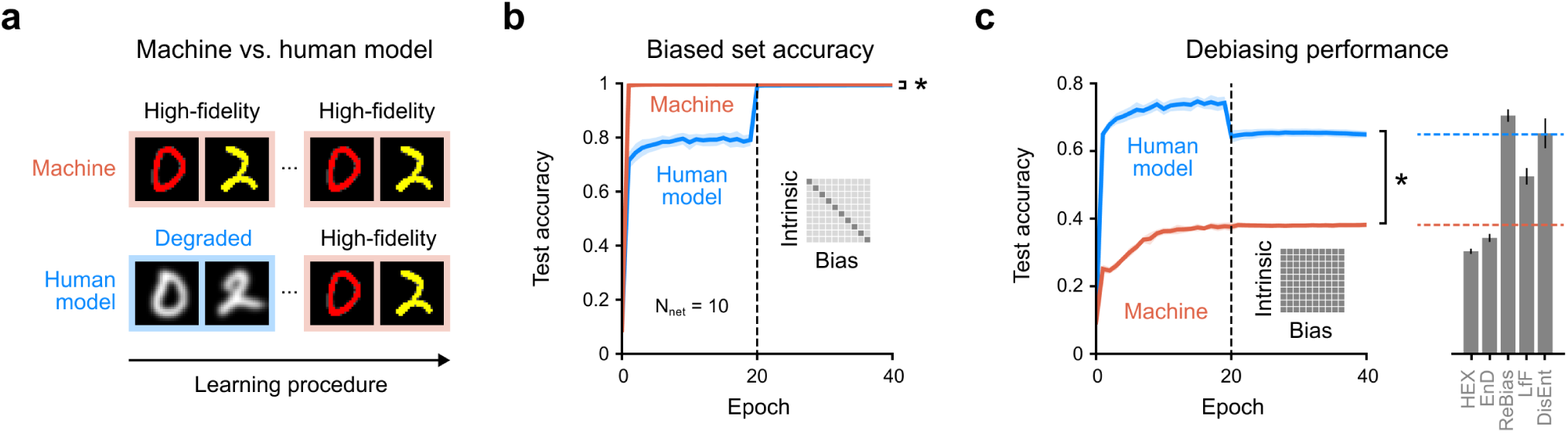
Debiased learning on the Color MNIST dataset through initial sensory degradation. (a) Training paradigms for conventional and human-inspired models. The conventional model is trained on high-fidelity images throughout, whereas the human model undergoes gradual sensory maturation — starting with degraded (blurry and grayscale) images, then transitioning to refined inputs. Both models are trained on a biased dataset in which intrinsic features (digit identity) are spuriously correlated with bias attributes (color). (b) Test accuracy on the biased test set that preserves the spurious correlations found in training. The dashed line at epoch 20 indicates the transition to high-fidelity inputs. (c) Debiasing performance on the unbiased test set with spurious correlations removed (left). Final test accuracy is compared to state-of-the-art debiasing methods (right) (HEX ^34^, EnD ^35^, ReBias ^36^, LfF ^31^, DisEnt ^25^). Dashed lines indicate the final performances of the conventional (orange) and human-inspired (blue) models.

**Supplementary Table. 1.** Benchmarked test accuracy on the unbiased test sets of Corrupted CIFAR-10 and Color MNIST. Test accuracy is reported across different levels of spurious correlation in the training data. A ratio of 99.5% denotes that 99.5% of training samples are bias-aligned and 0.5% are bias-conflict. Baseline machine and human model performances are given as mean *±* standard deviation over ten independent runs. Performance of existing debiasing algorithms (HEX ^34^, EnD ^35^, ReBias ^36^, LfF ^31^, DisEnt ^25^) is reproduced from prior work ^25^.

|  | Ratio (%) | Machine | Human Model | Debiasing Algorithms |  |  |  |  |
| --- | --- | --- | --- | --- | --- | --- | --- | --- |
|  |  |  |  | HEX | EnD | ReBias | LfF | DisEnt |
| Corrupted<br>CIFAR-10 | 99.5 | 18.36 ± 0.32 | 37.09 ± 0.44 | 13.87 ± 0.06 | 22.89 ± 0.27 | 22.27 ± 0.41 | 28.57 ± 1.30 | 29.95 ± 0.71 |
| | 99 | 23.45 ± 0.93 | 39.70 ± 0.55 | 14.81 ± 0.42 | 25.46 ± 0.41 | 25.72 ± 0.20 | 33.07 ± 0.77 | 36.49 ± 1.79 |
| | 98 | 30.91 ± 0.67 | 44.21 ± 0.46 | 15.20 ± 0.54 | 31.31 ± 0.35 | 31.66 ± 0.43 | 39.91 ± 0.30 | 41.78 ± 2.29 |
| | 95 | 45.06 ± 0.61 | 52.88 ± 0.35 | 16.04 ± 0.63 | 40.26 ± 0.85 | 43.43 ± 0.41 | 50.27 ± 1.56 | 51.13 ± 1.28 |
| Color<br>MNIST | 99.5 | 38.14 ± 0.69 | 64.89 ± 1.03 | 30.33 ± 0.76 | 34.28 ± 1.20 | 70.47 ± 1.84 | 52.50 ± 2.43 | 65.22 ± 4.41 |
| | 99 | 53.83 ± 0.43 | 75.30 ± 0.71 | 43.73 ± 5.50 | 49.50 ± 2.51 | 87.40 ± 0.78 | 61.89 ± 4.97 | 81.73 ± 2.34 |
| | 98 | 68.58 ± 0.65 | 83.61 ± 0.45 | 56.85 ± 2.58 | 68.45 ± 2.16 | 92.91 ± 0.15 | 71.03 ± 2.44 | 84.79 ± 0.95 |
| | 95 | 82.06 ± 0.65 | 89.90 ± 0.24 | 74.62 ± 3.20 | 81.15 ± 1.43 | 96.96 ± 0.04 | 80.57 ± 3.84 | 89.66 ± 1.09 |

**Supplementary Fig. 5.**
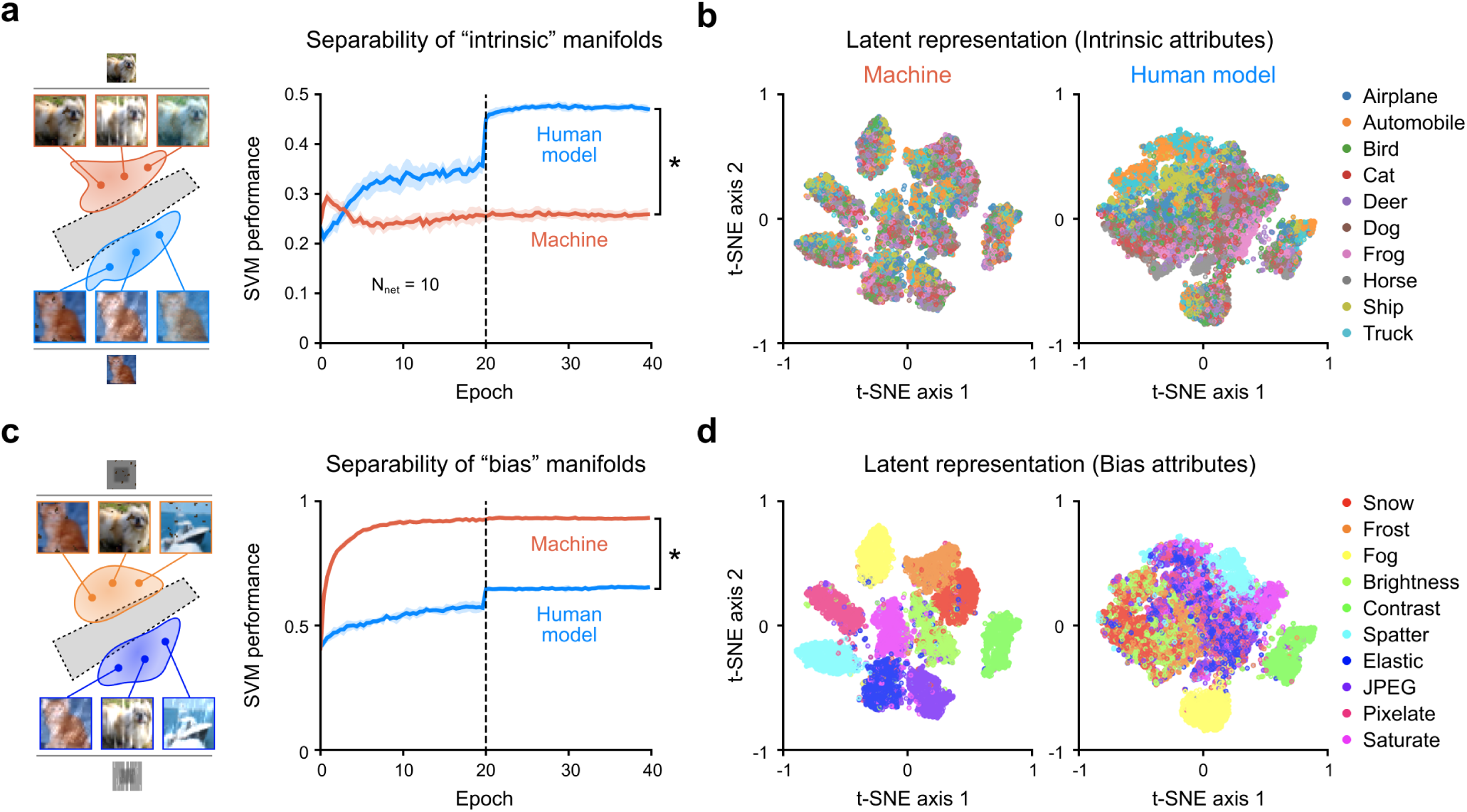
Latent representations of the machine and human model trained on Corrupted CIFAR-10. (a) Intrinsic manifold analysis. Latent feature vectors from an intermediate layer are grouped by intrinsic attributes (e.g., object or animal class). Separability is measured as the average classification accuracy of SVM classifiers trained on these features — higher scores indicate stronger encoding of intrinsic structure (human model vs. Machine, *n*_subject_ = 10, Wilcoxon rank-sum test, *P <* 10*^−^*^3^). The dashed line at epoch 50 denotes the onset of high-fidelity input exposure in the human model’s training. (b) Visualization of intrinsic structure. Intermediate-layer activations are projected into two dimensions using t-distributed stochastic neighbor embedding (t-SNE). Each point is colored according to its intrinsic class label. (c) Bias manifold analysis. Same analysis as in (a), but latent features are grouped by spurious bias attributes (e.g., corruption type), quantifying how much bias information is encoded (Human model vs. Machine, *n*_subject_ = 10, Wilcoxon rank-sum test, *P <* 10*^−^*^3^). (d) Visualization of bias structure. t-SNE projection as in (b), but colored by bias attributes instead of intrinsic class.

**Supplementary Fig. 6.**
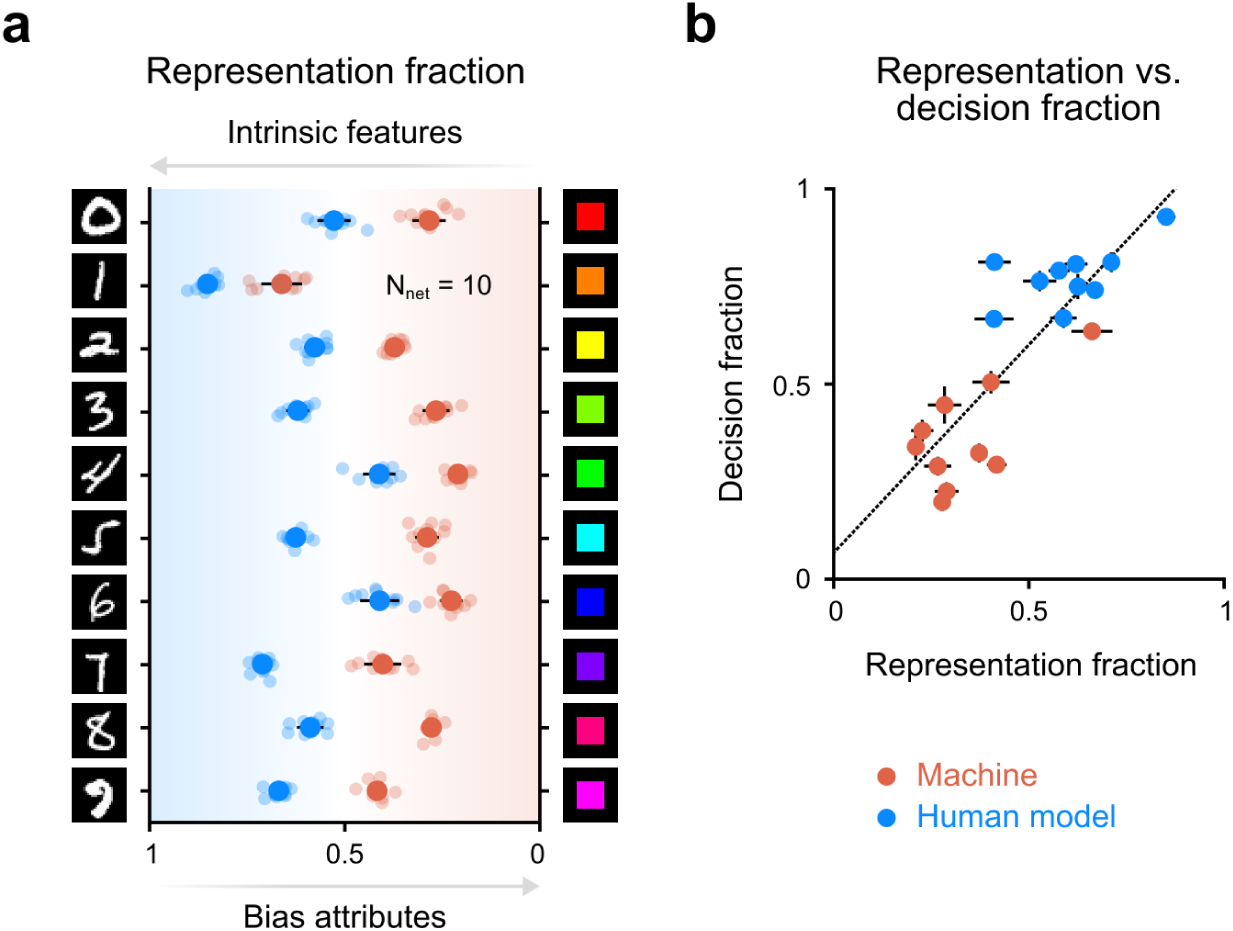
Debiased representations enable debiased classification. (a) Representation fraction (RF) quantifies the extent to which latent representations preferentially encode intrinsic attributes over bias attributes. An RF close to 0 indicates bias-dominated encoding, whereas a value near 1 indicates strong intrinsic encoding. Orange and blue circles represent machine and human model performances, respectively. (b) Relationship between representation and decision. The representation fraction is positively correlated with the decision fraction.

**Supplementary Fig. 7.**
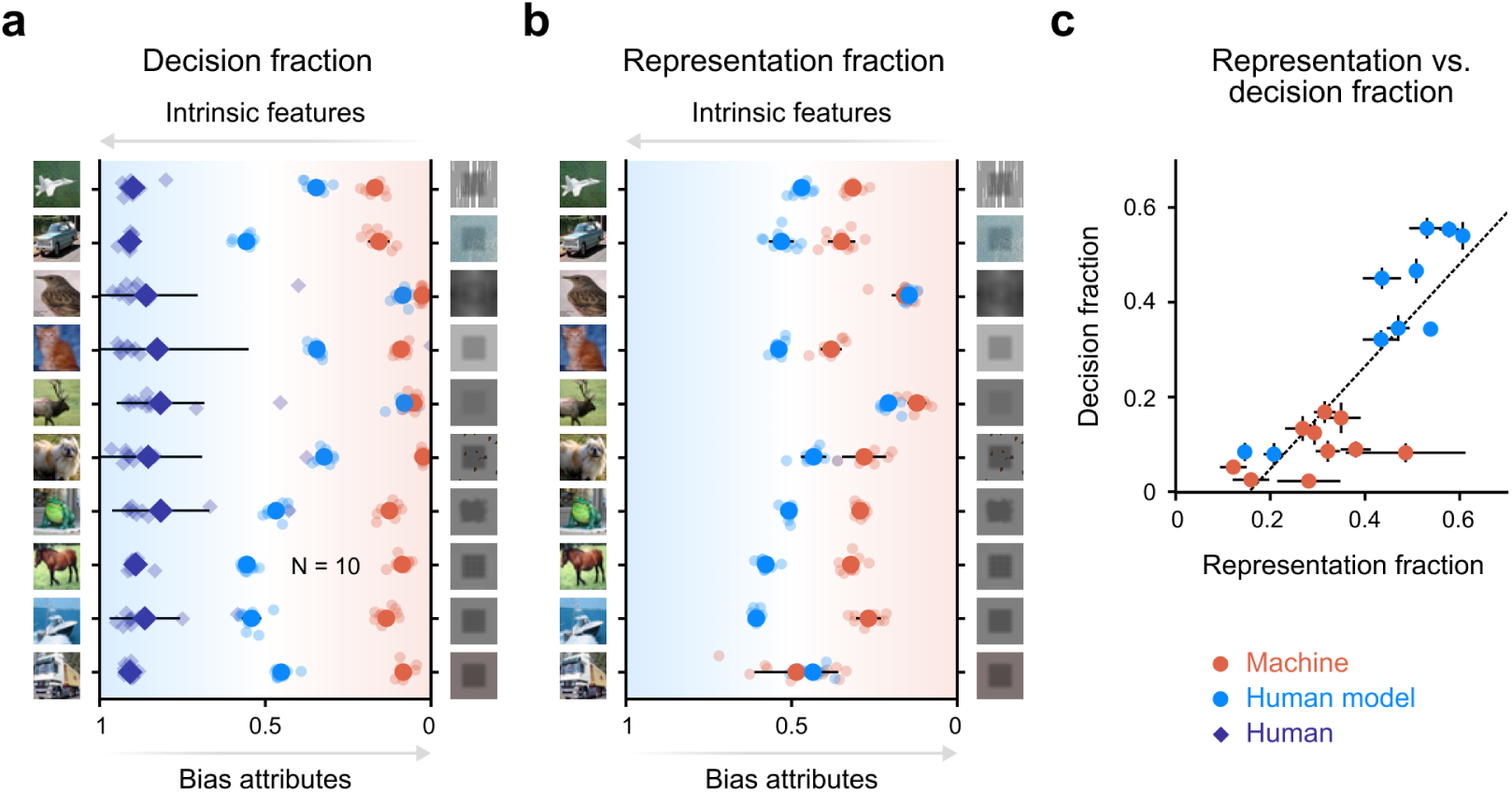
Debiased representations enable intrinsic-driven classification in corrupted CIFAR-10. (a) Decision fraction (DF) quantifies the extent to which model predictions rely on intrinsic features versus spurious bias. A DF near 0 indicates bias-driven decisions, while a value near 1 reflects intrinsic-feature-based decisions. Orange and blue circles indicate results from the conventional and human models, respectively. Purple diamonds represent human psychophysics data. (b) Representation fraction (RF) measures how strongly the model’s latent representations encode intrinsic versus bias attributes. Values closer to 1 indicate greater intrinsic encoding. (c) Relationship between representation and decision. A strong positive correlation is observed between RF and DF (regression line, *y* = 1.08*x −* 0.27, *R*^2^ = 0.698; Pearson correlation coefficient, *r* = 0.835, *P <* 10*^−^*^3^).

